# Long-read sequencing reveals transposable element-derived chimeric transcripts at zygotic genome activation in mammalian embryos

**DOI:** 10.64898/2026.05.25.727629

**Authors:** Saki Kawakami, Koichi Kitao, Shuntaro Ikeda, Shinnosuke Honda

## Abstract

**Background:** Transposable elements (TEs) are mobile genomic sequences that constitute one-third to one-half of the mammalian genome. Recently, TEs have been recognized for their important roles as cis-regulatory elements. TEs are broadly activated during zygotic genome activation (ZGA) in mammalian embryos, where they function as alternative promoters of host genes and drive the transcription of chimeric transcripts. However, the construction of comprehensive chimeric transcript databases based on short-read sequencing remains limited due to the repetitive and abundant nature of TEs in the genome. Here, we used long-read RNA sequencing to construct a comprehensive dataset of chimeric transcripts expressed in ZGA mouse and bovine embryos.

**Results:** We identified 11,996 and 4,755 chimeric transcripts variants derived from 2,695 and 1,200 host genes in mouse and bovine, respectively, exceeding the numbers reported in previous short-read-based studies. Among them, 114 orthologous pairs produced chimeric transcripts in both species. Gene Ontology analysis revealed significant enrichment of terms related to transcriptional regulation and protein modification in mouse, whereas no terms were significantly enriched in bovine. Assessment of the protein-coding potential of the TE-driven transcripts using predicted open reading frames (ORFs) revealed that the proportion of “Protein-coding” transcripts was lower, whereas that of “LncRNA” (long non-coding RNA) was higher compared with all transcripts in both species. Among the ORFs classified as “Protein-coding”, comparison with canonical ORFs revealed a tendency for the N terminus to be truncated while the C terminus remained intact in both species. TE-derived promoters used in mouse were enriched for mouse-specific TEs, whereas those in bovine were enriched for older TEs conserved among eutherians. In addition, long-read sequencing detected a greater number and proportion of TEs used as promoters in mouse and bovine than short-read sequencing. Although motif analysis identified KLF5 and OTX2 binding sites upstream of TE-derived promoters in both species, the specific TEs containing these motifs differed between the two species.

**Conclusions:** This study presents the first long-read sequencing analysis of chimeric transcripts in mammalian embryos in two species. Our approach revealed the functional similarities of chimeric transcripts between species, as well as species-specific differences in their TE compositions.

## Background

Transposable elements (TEs) are mobile genomic sequences that constitute approximately one-third to one-half of the mouse genome [1], [2], [3]. Based on their mode of transposition, TEs are broadly classified as class I and class II elements. Class I TEs propagate within the host genome via a “copy-and-paste” mechanism that involves an RNA intermediate [4]. Also referred to as retrotransposons, class I TEs in mammals are further divided into three major groups: long interspersed nuclear elements (LINEs), short interspersed nuclear elements (SINEs), and long terminal repeat (LTR) retrotransposons. Class II TEs are also known as DNA transposons and primarily move within the genome through a “cut-and-paste” mechanism in which their DNA sequences are excised and reinserted at new genomic locations [5].

Activation of TEs can lead to their insertion into the host genome, potentially disrupting essential genes or perturbing the regulation of nearby genes. In addition, TEs can produce mRNAs and proteins that are deleterious to the host. Consequently, hosts have evolved multiple mechanisms to silence TE expression, including epigenetic modifications, RNA degradation pathways, and KRAB domain-containing factors [6]. However, recent studies have demonstrated that hosts not only eliminate or suppress TEs as harmful elements but also co-opt them as regulatory components of gene expression[7]. During zygotic genome activation (ZGA) in mammalian embryos, TEs become broadly activated and contribute to gene regulation by serving as cis-regulatory elements, such as promoters, enhancers, and chromatin boundaries [8]. Notably, LTR retrotransposons, LINEs, and SINEs contain transcription start sites (TSSs) with intrinsic promoter activity and transcription factor-binding motifs [8]. Owing to these properties, TEs can act as alternative promoters and drive the transcription of downstream host genes in early embryos, producing TE-driven chimeric transcripts [9], [10], [11]. A subset of these chimeric transcripts possesses functions distinct from those of canonical mRNAs and proteins and influences early embryonic development in mouse [11], [12].

Although the importance of chimeric transcripts in mammalian embryos is being recognized, a comprehensive and accurate database of chimeric transcripts has yet to be established, largely due to the technical limitations of conventional RNA sequencing. Standard short-read RNA sequencing determines sequences by reading short fragments, typically 50–300 nucleotides. While this method offers high base-level accuracy, it cannot capture full-length mRNA. In particular, accurate mapping of TE-driven transcripts remains challenging for short-read sequencing because TEs have repetitive sequences and are abundant throughout the genome. On the other hand, long-read sequencing enables the sequencing of mRNA molecules up to several kilobases long as a single contiguous sequence. Consequently, many studies on TEs have highlighted the limitations of short-read sequencing and emphasized the need for full-length mRNA analysis [12], [13].

In this study, we performed long-read RNA sequencing to construct a comprehensive catalog of TE-driven chimeric transcripts expressed in mammalian preimplantation embryos. Our results suggest the advantages of long-read sequencing for chimeric transcript analysis and highlight the species-specific nature of TE-driven chimeric transcripts.

## Methods

### In vitro fertilization and embryo culture in mouse

Eight- to eleven-week-old female Slc:ICR mouse (Japan SLC, Hamamatsu, Japan) were superovulated by injection of 7.5 IU of equine chorionic gonadotropin (ASKA Pharmaceutical, Tokyo, Japan), followed 48 h later by 7.5 IU of human chorionic gonadotropin (ASKA Pharmaceutical). Cumulus-oocyte complexes were harvested 14 h after human chorionic gonadotropin injection and placed in human tubal fluid (HTF) medium [14] supplemented with 4 mg/mL bovine serum albumin (A3311; Sigma-Aldrich, St. Louis, MO) under paraffin oil (Nacalai Tesque, Kyoto, Japan). Spermatozoa were collected from the cauda epididymis of 13–20-week-old male Slc:ICR mouse (Japan SLC) and capacitated by culturing in HTF medium for 1 h. After in vitro fertilization (IVF), embryos were washed with HTF medium to remove cumulus cells. The embryos were then transferred to potassium simplex optimized medium (KSOM) [15] containing 1 mg/mL bovine serum albumin and amino acids and cultured at 37°C in 5% CO_2_ until 36 h after fertilization. Embryos that reached the late two-cell stage were then collected for RNA extraction.

### In vitro fertilization and embryo culture in bovine

In vitro production of bovine embryos by IVF and embryo culture was performed as previously described [16]. Bovine ovaries used in this study were purchased from a commercial abattoir as by-products of meat processing, and the frozen bull semen used for IVF was also commercially available. In brief, bovine oocytes were recovered from the ovaries of Japanese Black or Japanese Black × Holstein F1 bovine. Cumulus-enclosed oocytes were in vitro-matured for 22 h in Medium 199 with Earle’s salts (Life Technologies) supplemented with 5% (v/v) fetal calf serum and 0.2 IU/mL follicle-stimulating hormone (Kyoritsu Seiyaku). Matured cumulus-enclosed oocytes were subjected to IVF with sperm from a Japanese Black bull. At 20 h post-insemination, one-cell embryos were freed from cumulus cells and then in vitro-cultured in in vitro culture medium. The cultures were performed at 38.5°C in 5% CO_2_/air (in vitro-matured and in vitro-fertilized) or in 5% CO_2_, 5% O_2_, and 90% N_2_ (in vitro-cultured). At 72 h post-insemination, 8-cell to 16-cell stage embryos were collected for RNA extraction.

### RNA extraction

Embryos were washed in 50 µL of phosphate-buffered saline containing 0.5% polyvinylpyrrolidone (PVP K-30; Nacalai Tesque), dissolved in TRIzol reagent (Thermo Fisher Scientific Inc., Waltham, MA) at a concentration of 1 µL/embryo, vortexed for 1 min, and stored at −80°C. After thawing, the TRIzol reagent containing the extracted RNA was brought up to a total volume of 300 µL to pool RNA from approximately 200 embryos. RNA extraction was performed following the manufacturer’s protocol. During isopropanol precipitation, 1 µL of glycogen (F. Hoffmann-La Roche Ltd., Basel, Switzerland) was added to facilitate co-precipitation. The air-dried pellet was dissolved in 10 µL of UltraPure Distilled Water (Thermo Fisher Scientific Inc.) and the concentration was measured with a Qubit RNA HS Assay Kit (Thermo Fisher Scientific Inc.).

### Long-read cDNA library preparation and nanopore sequencing

The cDNA-PCR Sequencing Kit (SQK-PCS114; Oxford Nanopore Technologies Plc., Oxford, UK) was used for reverse transcription and library preparation according to the manufacturer’s protocol. After reverse transcription, cDNA was amplified using LongAmp Hot Start Taq 2X Master Mix (New England Biolabs) with 14 cycles. Amplified cDNA was purified using AMPure XP beads (Beckman Coulter, Inc., Brea, CA) and quantified using a Qubit 1X dsDNA HS Assay Kit (Thermo Fisher Scientific Inc.). Each library was separately loaded into an R10.4.1 flow cell (Oxford Nanopore Technologies Plc.) and sequenced on a MinION sequencer (Oxford Nanopore Technologies Plc.).

### Processing of long-read RNA-seq data

Mouse and bovine Fast5 files were basecalled using MinKNOW (v6.0.14) or Dorado (v0.6.3 or v0.7.3) to generate FASTQ files. The FASTQ reads were then processed and mapped to reference genomes using the wf-transcriptomes pipeline in EPI2ME (v1.7.1) (https://github.com/epi2me-labs/wf-transcriptomes) to obtain GTF files. The mouse reference genome (mm10) and annotation data were downloaded from Illumina iGenomes (http://support.illumina.com/sequencing/sequencing_software/igenome.ilmn), while the bovine reference genome and annotation data were downloaded from the UCSC Genome Browser database (https://hgdownload.soe.ucsc.edu/goldenPath/bosTau9/bigZips/).

### Identification and classification of TEs used as promoters

The TSS was defined as the first nucleotide at the start position of each detected transcript. For filtering, transcripts < 200 bp in length were eliminated from the GTF files generated by Epi2ME. Then, the TSS of each transcript was extracted from the GTF file and converted into a BED file. RepeatMasker annotations of TEs were downloaded from the UCSC Genome Browser database (mouse: https://hgdownload.soe.ucsc.edu/goldenPath/mm10/bigZips/mm10.fa.out.gz; cow: https://hgdownload.soe.ucsc.edu/goldenPath/bosTau9/bigZips/bosTau9.fa.out.gz).

Using bedtools intersect (BEDTools v2.31.1; https://bedtools.readthedocs.io/en/latest/) [17], TSSs that overlapped TEs (TE-TSSs) were extracted. To eliminate redundancy among transcript variants, duplicate entries were removed using a custom script. Subsequently, the TE-TSSs were intersected (BEDTools, v2.31.1) and quantified based on their class and subfamily using another custom script.

### Comparison of TE-TSS detection between long-read and short-read sequencing

Short-read RNA-seq data were downloaded from GSE71434 for mouse late two-cell embryos and from GSE52415 for bovine eight-cell embryos. The FASTQ files were mapped using STAR (v2.7.10b) with the “--outSAMtype BAM SortedByCoordinate” and “--quantMode TranscriptomeSAM” options to obtain BAM files [18]. The BAM files were assembled into GTF files with StringTie2 (v2.2.1) using the “--conservative” option and the same reference annotation (“-G” option) used for long-read RNA-seq processing [19]. The GTF files from biological replicates were merged for each species using StringTie2 (v2.2.1). The TSSs of each transcript were extracted, and duplicate entries were removed as described above. To account for differences in sequencing platforms and potential errors in TSS identification, TSSs from the short-read datasets were extended 50 bp upstream and downstream using bedtools slop (BEDTools, v2.31.1). Long-read TSSs that did not overlap any extended short-read TSSs were defined as “Long-read only” TSSs, while those that did overlap were defined as “Shared with short-read” TSSs. The same analysis was performed under the opposite condition (short-read TSSs filtered by long-read TSSs) to obtain “Short-read only” TSSs and “Shared with Long-read” TSSs. Finally, TE-TSSs in long-read and short-read datasets were analyzed as described above.

### Identification of host genes associated with TE-driven chimeric transcripts

After removing reads shorter than 200 bp, transcripts classified by GffCompare (in EPI2ME) with the following class codes were filtered out from subsequent chimeric transcript analysis: *s* (intron match on the opposite strand), *x* (exonic overlap on the opposite strand), *I* (fully contained within a reference intron), *y* (contains a reference with its introns), *p* (possible polymerase run-on), *r* (repeat), or *u* (none of the above). The “transcript_id” values from BED files containing all TSSs overlapping TEs were searched in the GTF file generated in the “Processing of long-read RNA-seq data” section to extract the corresponding transcript entries of candidate chimeric transcripts using a custom script.

### Acquisition of chimeric transcript data from previous studies

The study by Modzelewski et al. was based on two mouse RNA-seq datasets obtained by Deng et al. and Xue et al [12], [20], [21]. Of these two datasets, we chose the dataset from Deng et al. because it was published more recently. Then, gene symbols whose entries contained either “promoter” or “promoter_tss” in the “type” column of Table S3 were extracted. From the data of Oomen et al., gene symbols whose entries contained “late-2-cell” in the “stages” column of Table S5 were extracted. Duplicate entries were removed using a custom script, and the final set of gene symbols was used to represent the TE-driven genes detected in the previous studies.

### Identification of conserved chimeric transcripts between mouse and bovine

Candidate chimeric transcripts identified in both mouse and bovine were compared to extract those conserved between the two species. Gene symbols of candidate chimeric transcripts in both species were converted into Ensembl gene IDs, and common orthologs identified as candidate chimeric transcripts were extracted based on the “btaurus_homolog_ensembl_gene” attribute of the mm10 dataset in Ensembl BioMart (https://www.ensembl.org/info/data/biomart/index.html) using the R biomaRt package (v2.64.0).

### Gene Ontology analysis

Gene Ontology (GO) analysis was conducted for the molecular function and biological process categories using the clusterProfiler R package (v4.16.0). Enriched GO terms were identified using *p*-value and *q*-value cutoffs of < 0.05.

### Prediction of the protein-coding potential of transcripts

Transcript sequences were extracted from GTF files and converted into FASTA format using gffread (v0.12.7) [22]. The protein-coding potential of each transcript was then predicted using CPAT (v3.0.5) [23]. For mouse, the prebuilt hexamer table and logistic regression model provided by CPAT were used. For bovine, species-specific models were constructed from 10,000 randomly selected entries from ncbiRefSeq.gtf, which was downloaded from the UCSC Genome Browser database (https://hgdownload.soe.ucsc.edu/goldenPath/bosTau9/bigZips), and 10,000 randomly selected entries from ncrna.fa, which was downloaded from the Ensembl database (https://ftp.ensembl.org/pub/release-115/fasta/bos_taurus/). The “--top-orf=1” option was applied to extract the longest ORF. The default CPAT cutoff value (0.44) was used for mouse, while the cutoff value for bovine (0.38) was determined based on a receiver operating characteristic curve generated using the data employed for model construction. Transcripts with coding probability values above the cutoff were classified as “Protein-coding”. Transcripts with coding probability values equal to or below the cutoff were classified as “LncRNA” if their length was greater than 200 bp; all others were classified as “Else”.

### Identification of novel proteins and comparison with reference protein sequences

From a list of transcript IDs assigned as protein-coding genes, the corresponding transcript sequences were obtained using seqkit grep with the -f option [24]. Next, the longest putative ORF for each transcript was extracted using TransDecoder.Predict in TransDecoder (v5.7.1) with the “-G universal -m 100 --single_best_only” options [25]. The predicted ORF sequences were searched against reference protein databases using the blastp command in BLAST+ (v2.17.0) with the “-max_target_seqs 1 -evalue 1e-5” options, and results were obtained in outfmt 6 format [26]. Reference protein sequences for both mouse (*Mus musculus*) and bovine (*Bos taurus*) were downloaded from UniProtKB (https://www.uniprot.org/uniprotkb; accessed November 27, 2025).

To identify transcripts harboring ORF sequences that confidently matched at least a portion of their subject sequences (i.e., reference protein sequences), only ORFs with an alignment length ≥ 50% of the subject length and a sequence identity ≥ 95% to the subject sequences were retained. After filtering, the lengths of the predicted ORFs were compared with the corresponding subject lengths. Using the values of qstart, qend, sstart, send, qlen, and slen obtained from the outfmt 6 output, four metrics representing the relative deletion or extension at each terminus were calculated as follows:

N_Del = (sstart − 1) / slen (Relative length of the unaligned N-terminal region in the subject sequence)

C_Del = (slen − send) / slen (Relative length of the unaligned C-terminal region in the subject sequence)

N_Ext = (qstart − 1) / slen (Relative length of the unaligned N-terminal region in the query sequence)

C_Ext = (qlen − qend) / slen (Relative length of the unaligned C-terminal region in the query sequence)

Each terminus (N- and C-) was classified as “Intact”, “Del“, “Ext”, and “Alt” based on the following criteria: Intact = N(C)_Del < 0.1 and N(C)_Ext < 0.1; Del = N(C)_Del ≥ 0.1 and N(C)_Ext < 0.1; Ext = N(C)_Del < 0.1 and N(C)_Ext ≥ 0.1; Alt = N(C)_Del ≥ 0.1 and N(C)_Ext ≥ 0.1. Transcripts that were excluded by filtering, as well as those without any hit against the reference database, were classified as “Others”.

### Investigation of the conservation of transcripts classified as “Others”

The conservation of ORFs classified as “Others” was investigated using hmmscan from HMMER (v3.4) against the Pfam protein domain database (v38.1) and using BLASTP [27], [28] against the UniProtKB database. When multiple hits were obtained for a single ORF sequence, the hit with the lowest *E*-value was selected for both searches.

### Motif analysis of TEs used as promoters

Known motifs were searched for in the 1,000-bp upstream sequences of all TSSs and TE-TSSs. Motif enrichment analysis was performed using findMotifsGenome.pl in HOMER (v5.1) with the “-mset vertebrate” and “-size given” options [29]. To perform motif enrichment analysis with background sequences, the 1,000-bp upstream sequences of all TSSs were used as background sequences with the “-bg” option. TE subfamilies containing each enriched motif were identified using scanMotifGenomeWide.pl in HOMER (v5.1). For both mouse and bovine, sequences of TE subfamilies identified as TE-TSSs were downloaded from the same RepeatMasker files used in the “Identification and classification of TEs used as promoter” section. Corresponding FASTA files were then generated using bedtools getfasta (BEDTools, v2.31.1). After scanning subfamily sequences for the presence of motifs, the number of motif hits was counted. The ratio of motif-containing elements was then calculated by dividing the number of hits by the total number of entries in the corresponding BED file for each subfamily.

## Results

### Long-read sequencing may provide an advantage in detecting chimeric transcripts

Long-read RNA sequencing obtained 1,562,691 reads from late 2-cell stage mouse embryos and 2,122,809 reads from 8- to 16-cell stage bovine embryos (Supplementary Table 1). In total, 11,996 transcripts from mouse and 4,755 transcripts from bovine were identified as chimeric transcripts in this study (see workflow in Fig. 1). After removing redundancy among splicing variants, we identified 2,695 genes in mouse and 1,200 in bovine as TE-driven genes, including some not previously identified in public short-read sequencing data [21] (Fig. 2A, 2B). Subsequently, we compared the TE-driven genes identified by long-read sequencing with those from two previous studies that used short-read sequencing data of mouse ZGA-stage embryos [12], [13]; 588 genes (42.6% of 1,380) were common with the first study, while 561 genes (36.1% of 1,555) were common with the second study (Fig. 2C, 2D). In contrast, only 279 genes were identified as TE-driven genes in both previous studies (Fig. 2E). Among the genes identified in our long-read data, 114 orthologs were shared between mouse and bovine.

**Fig. 1.**
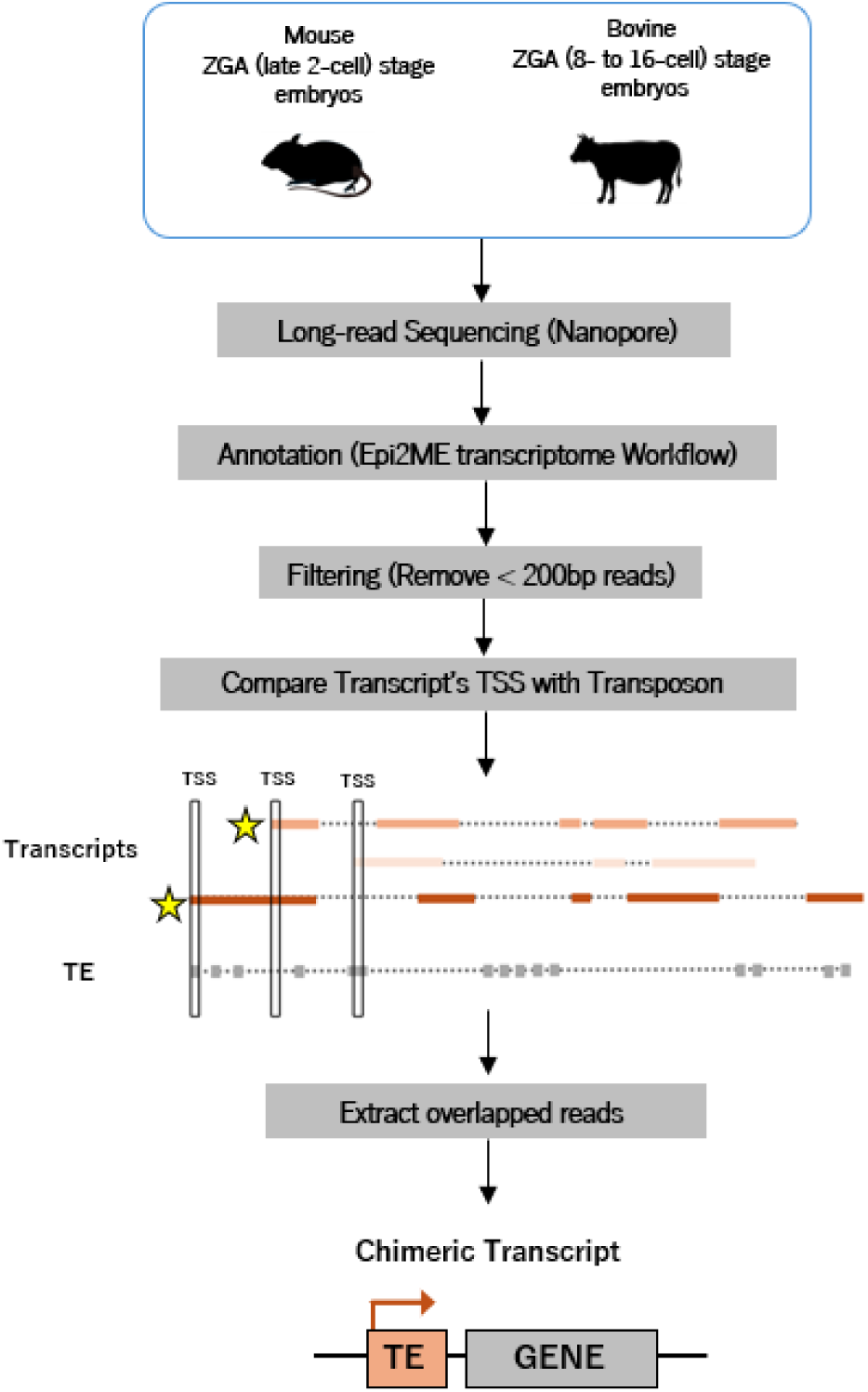
Workflow for the exploration of chimeric transcripts using long-read sequencing data. FASTQ data were processed using the wf-transcriptomes pipeline in Epi2ME (v1.7.1). The transcripts were mapped using reference genome and GTF annotation files obtained from Illumina iGenomes for mouse and the UCSC Genome Browser database for bovine. After removing short transcripts (< 200 bp), TSSs that overlapped TEs were extracted.

**Fig. 2.**
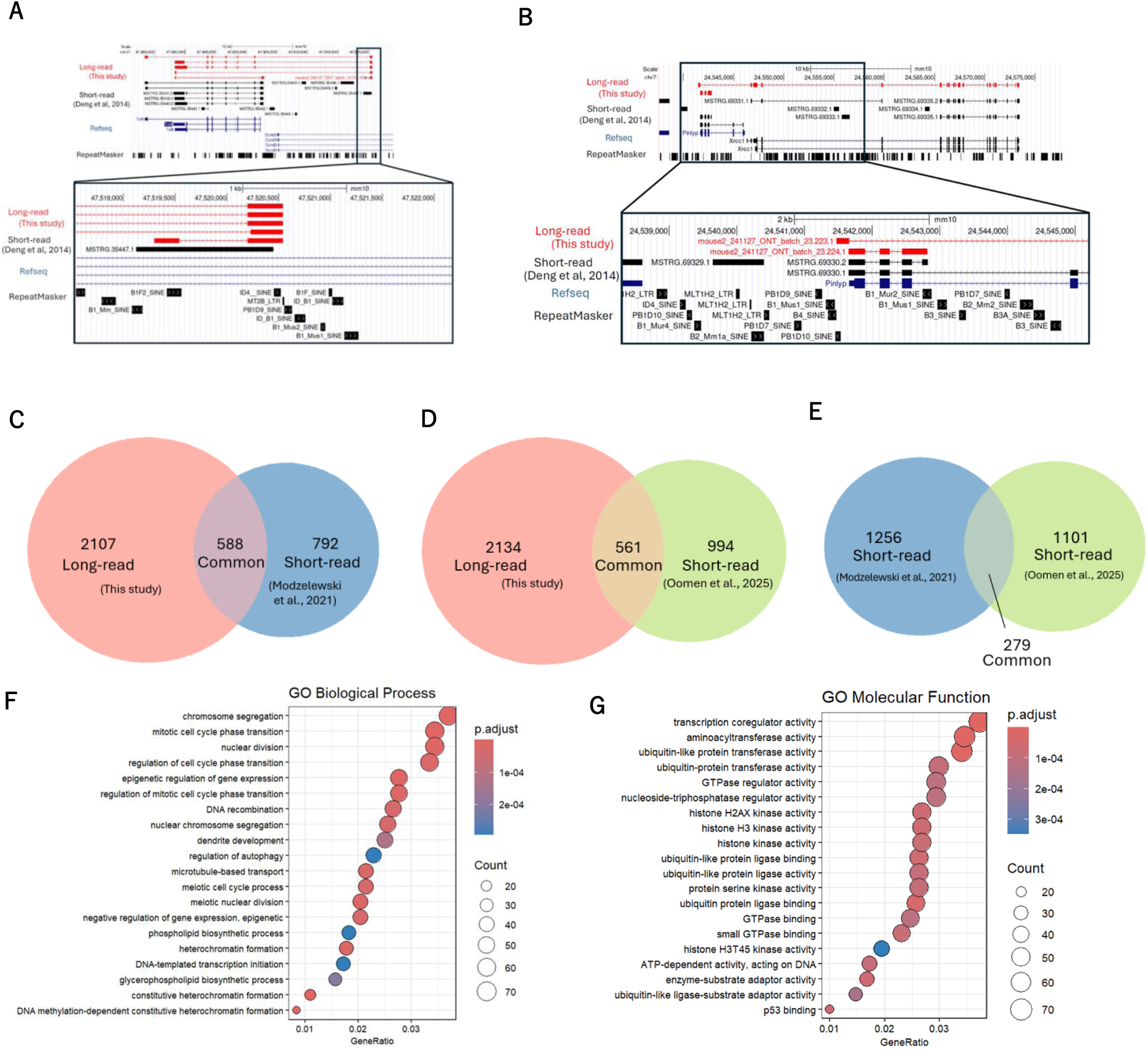
Comparison and functional characterization of chimeric transcripts detected in long- and short-read sequencing. (A, B) Representative IGV snapshots illustrating the genomic loci of (A) *Taf8* (TATA-box binding protein-associated factor 8) and (B) *Xrcc1* (X-ray repair cross-complementing protein 1) in the mouse genome (mm10). The upper panels show the overall gene structure, while the lower panels provide magnified views of the TSS regions. The TSSs of the chimeric transcripts overlap the MT2B (LTR) and PB1D10 (SINE) elements, respectively. (C–E) Venn diagrams showing the overlap of genes identified as chimeric transcripts in mouse between (C) this study and Modzelewski et al., (D) this study and Oomen et al., and (E) Modzelewski et al. and Oomen et al. (F, G) Gene Ontology enrichment analysis for genes identified as chimeric transcripts in mouse. The top 20 enriched terms for (F) Biological Process and (G) Molecular Function are shown. Statistical significance was defined as *p.adj* < 0.05 and *q* < 0.05.

### Functions of chimeric transcripts may differ between species

To investigate the biological functions associated with TE-driven genes, we performed GO enrichment analysis. In mouse, terms related to regulation of transcription and chromatin dynamics, such as “chromosome segregation”, “mitotic cell cycle phase transition”, and “nuclear division” were enriched in the biological process category (*p.adj* < 0.05, *q* < 0.05) (Fig. 2F). In the molecular function category, terms related to transcriptional regulation and protein modification were enriched, such as “transcription coregulator activity”, “aminoacyl transferase activity”, “ubiquitin-like protein transferase activity”, and “histone H3/H2AX kinase activity” (*p.adj* < 0.05, *q* < 0.05) (Fig. 2G). Conversely, none of the terms were significantly enriched in either the biological process or molecular function category in bovine (Supplementary Table 2A and 2B). Among the 114 orthologous genes shared between mouse and bovine, “DNA recombination” was enriched in the biological process category, and terms related to kinase activity, such as “histone H3/H2AX kinase activity” and “protein serine kinase activity”, were enriched in the molecular function category (*p.adj* < 0.05, *q* < 0.05) (Supplementary Table 2C and 2D).

### Chimeric transcripts function as lncRNAs and tend to produce translation products with N-terminal truncations

Protein sequences translated from chimeric transcripts may differ in function from those of their canonical counterparts [12], [11]. To investigate the protein-coding potential, we analyzed ORF sequences of TE-driven transcripts and classified them as “Protein-coding”, “LncRNA”, and “Else” (transcripts not belonging to either category, such as processed pseudogenes, snoRNAs, and rRNAs). In mouse, 3,448 TE-driven transcripts (30.3%) were predicted to be “Protein-coding”, 3,106 (27.3%) to be “LncRNA”, and 4,813 (42.3%) to be “Else” (Fig. 3A). In bovine, the corresponding numbers were 1,163 (25.2%), 1,953 (42.4%), and 1,492 (32.4%), respectively (Fig. 3B). In contrast, when all detected transcripts were considered, more than half of the transcripts were classified as “Protein-coding” in both species (mouse, 25,977 [63.8%]; bovine, 25,731 [64.3%]) (Fig. 3A, 3B). The numbers of “LncRNA” and “Else” were 6,354 (15.6%) and 8,393 (20.6%) in mouse and 9,056 (22.6%) and 5,204 (13.0%) in bovine, respectively (Fig. 3A, 3B).

**Fig. 3.**
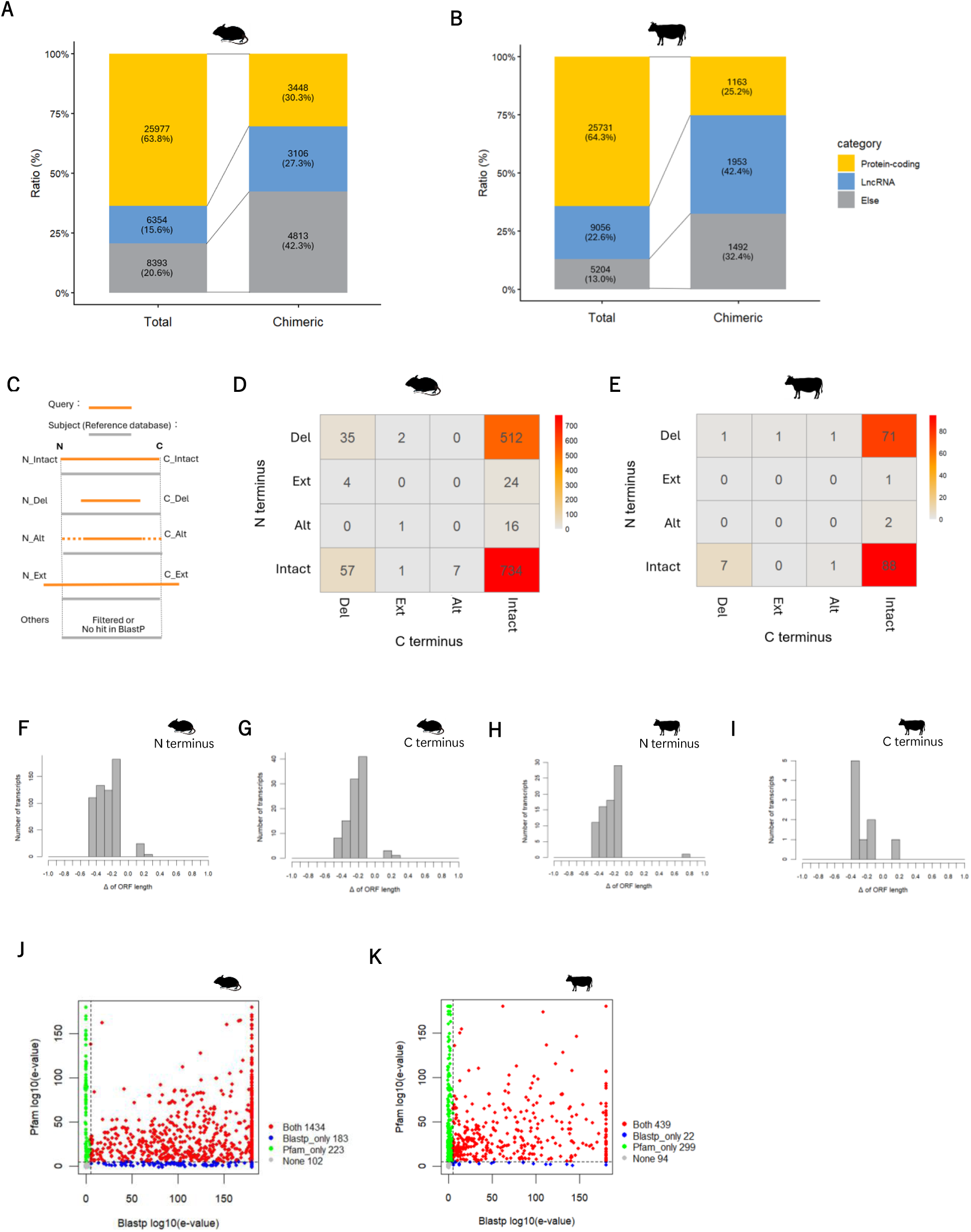
Protein-coding potential and ORF structure of chimeric mRNAs. (A, B) Classification of total and chimeric transcripts identified in (A) mouse and (B) bovine embryos according to their protein-coding potential (assessed by CPAT) and transcript length. (C) Schematic representation of the ORF structural categories: Intact, Del (deletion), Ext (extension), Alt (alternative), and Others. (D, E) Heatmap showing the distribution of ORF categories for both N and C termini among chimeric transcripts in (D) mouse and (E) bovine. Numbers within each box represent the count of transcripts. Only ORFs with hits against the UniProtKB database (via BLASTP) are shown. (F–I) Histograms representing the ratios of (F, H) N-terminal and (G, I) C-terminal deletions and extensions in ORFs of chimeric transcripts in (F, G) mouse and (H, I) bovine. ORFs classified as “Del” or “Ext” were analyzed relative to the lengths of the corresponding subject ORFs. (J, K) Scatter plots showing the BLASTP scores (x-axis) and Pfam scores (y-axis) for ORFs of chimeric transcripts classified as “others” in (J) mouse and (K) bovine. Significant hits were defined by an *E*-value threshold of 10^−5^ in both searches. Blue points indicate ORFs exceeding the BLASTP threshold only; green points indicate ORFs exceeding the Pfam threshold only; red points indicate ORFs meeting both criteria; and gray points indicate ORFs below both thresholds.

Next, we compared the predicted ORF sequences of “Protein-coding” chimeric transcripts with their canonical ORF sequences obtained from UniProtKB and evaluated differences at the N and C termini. In both mouse and bovine, the most abundant category comprised ORFs with N and C termini identical to their reference sequences (N_Intact, C_Intact), followed by ORFs with N-terminal deletions but intact C termini (N_Del, C_Intact) (Fig. 3C–E). We then examined the extent of N- and C-terminal changes in ORFs harboring “Del” or “Ext” relative to the reference ORF sequences. In both species, ORFs with “Del” were more frequently observed than those with “Ext” at both the N and C termini (Fig. 3F–I). In addition, among ORFs harboring “Del”, the number of ORFs decreased as the extent of the deletion increased, except at the C terminus in bovine (Fig. 3F–I).

Furthermore, 1,942 mouse and 854 bovine ORFs were classified as “Others”, comprising ORFs removed during filtering or absent from the UniProtKB database. To evaluate whether these ORFs encode functional proteins or contain known domains, we searched the UniProtKB and Pfam databases, respectively. Many ORFs classified as “Others” were predicted to encode proteins or to contain functional domains (Fig. 3J, 3K), suggesting that a substantial fraction retains biological function.

### Long-read sequencing detects TE-TSSs slightly more efficiently than short-read sequencing in both number and proportion

The number of TE-TSSs detected by long-read sequencing exceeded those identified by short-read sequencing in both mouse and bovine (Fig. 4A, 4B). In addition, the ratio of TE-TSSs to total TSSs was slightly higher in long-read than short-read sequencing: 27.8% (8,476 of 30,474) vs. 22.0% (4,913 of 22,346) in mouse, and 11.2% (3,253 of 29,040) vs. 9.5% (2,726 of 28,766) in bovine (Fig. 4A, 4B).

**Fig. 4.**
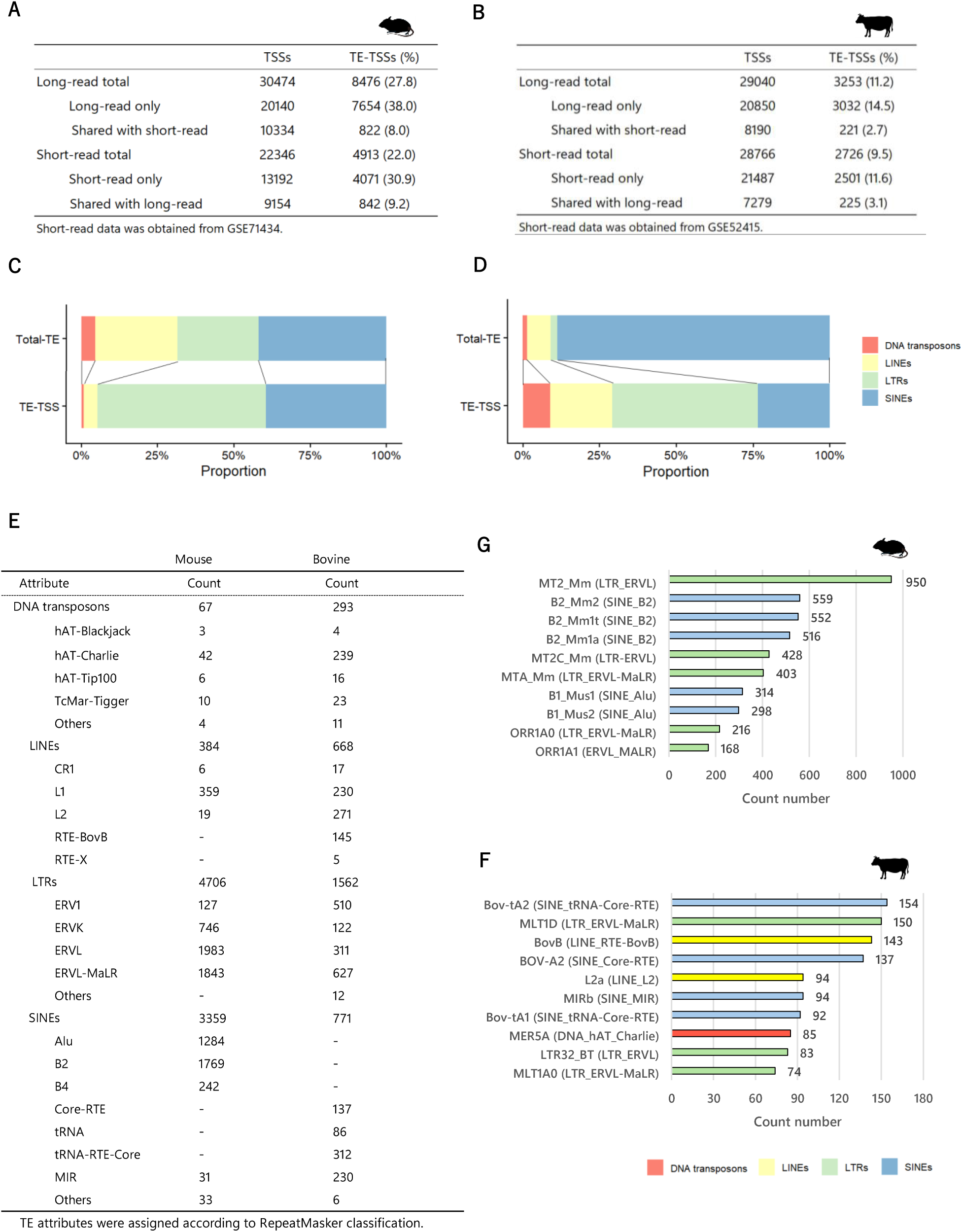
Genomic distribution and classification of TE-TSSs. (A, B) Comparison of the total number of TSSs and TE-TSSs detected by long- and short-read sequencing in (A) mouse and (B) bovine. (C, D) Comparison of the proportions of TEs acting as TE-TSSs, categorized by TE class and order, relative to the whole-genome distribution of TEs in (C) mouse and (D) bovine. (E, F) Number of TE-TSSs identified within the top 10 most frequent TE subfamilies (E) in mouse and (F) bovine. (G) Distribution of TE-TSS counts across TE classes, orders, and families in mouse and bovine.

### TEs used as promoters exhibit both shared and species-specific features

A total of 10,334 TSSs in mouse and 8,910 TSSs in bovine were identified by both long- and short-read sequencing. Although more than 20,000 TSSs were identified in both species and both platforms, fewer than half of the TSSs were shared between the two sequencing platforms in either species. This overlap ratio was even lower for TE-TSSs, with fewer than 10% of TE-TSSs commonly identified by both long- and short-read sequencing in either species.

Next, we categorized the TE classes of the TE-TSSs to examine commonalities and species-specific patterns. LTR retrotransposons were enriched in the TSS region in both species, whereas DNA transposons and LINEs were enriched in TSSs only in bovine (Fig. 4C, 4D). At the superfamily level, ERVL and ERVL-MaLR were enriched in TSSs in both mouse and bovine (Fig. 4E). In mouse, B2, a SINE superfamily member absents from the bovine genome, was highly ranked among TE-TSSs, whereas, in bovine, ERV1, an LTR retrotransposon rarely found in the mouse genome, was highly ranked (Fig. 4E). At the subfamily level, mouse-specific TEs such as MT2_Mm and MT2_B2 were highly ranked in mouse. In contrast, older TEs conserved across eutherian mammals, such as MLT1D and L2a, were highly ranked in bovine (Fig. 4F, 4G).

### Motif analysis at TE-TSSs shows low cross-species conservation

Motif analysis of the upstream sequences of all TSSs using a semi-random background revealed that SP1, RONIN, and KLF17 motifs were commonly enriched among the top 15 ranked motifs in both mouse and bovine (Supplementary Fig. 1A, 1B). On the other hand, only OTX2 and KLF5 motifs were commonly identified in the upstream sequences of TE-TSSs (Fig. 5A, 5B). Among the identified motifs, GATA3 and RAGR in mouse and DUXBL, DUX4, and YY1 in bovine were observed as motifs of known transcription factors functioning in embryos [30], [31], [32], [33], [34]. Furthermore, motif analysis of the upstream sequences of TE-TSSs was performed using the upstream sequences of all TSSs as background. In mouse, motifs such as KLF5, OTX2, GATA3, and ZNF134 were enriched, whereas, in bovine, motifs including DUXBL, BRN1, OCT4, and TEAD4 were enriched. Notably, no common motifs were observed among the top 15 enriched motifs in the two species (Supplementary Fig. 1C, 1D).

**Fig. 5.**
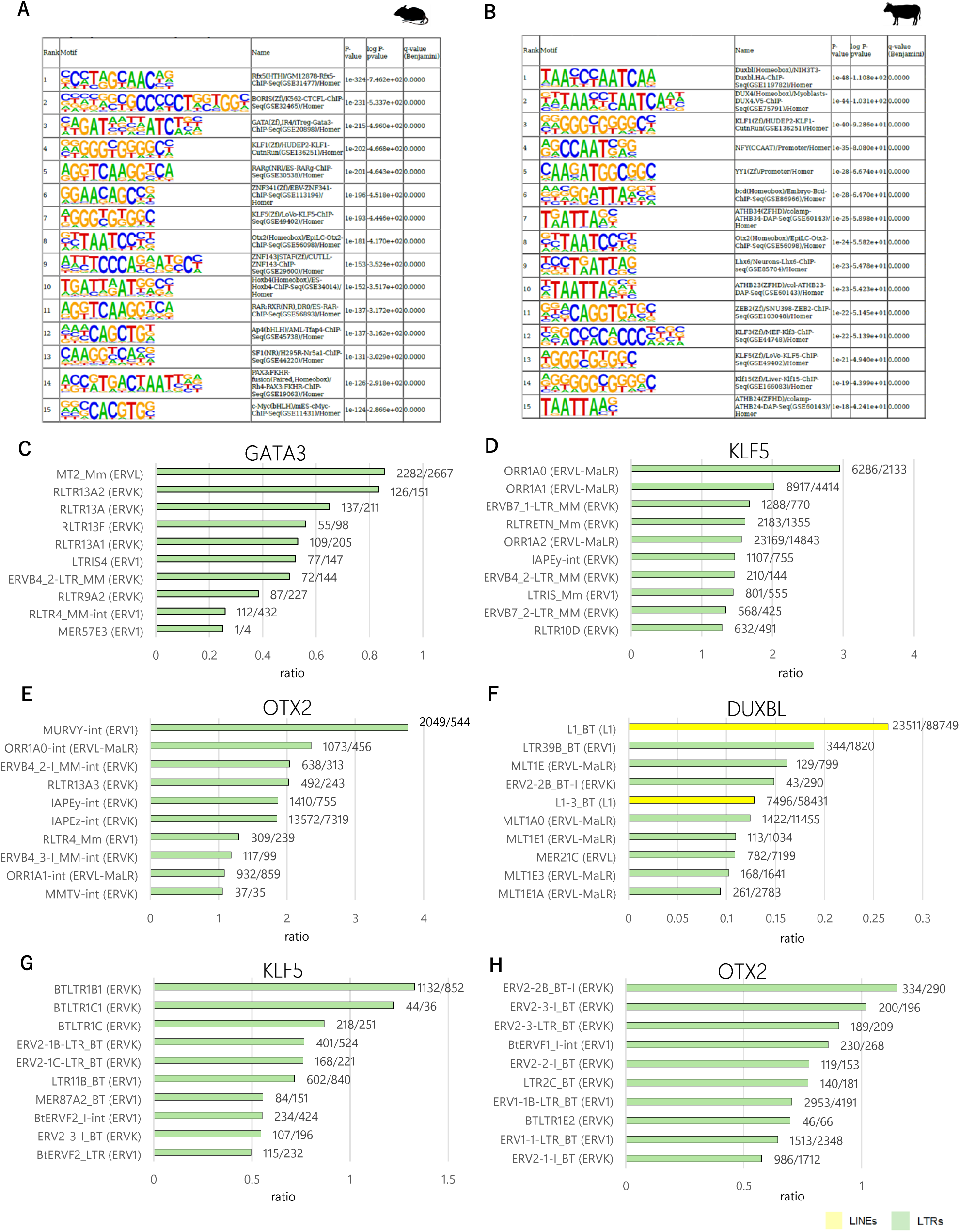
Motif analysis of the upstream regions of TE-TSSs. (A–B) The 15 most significantly enriched known motifs identified in regions within 1000 bp upstream of TE-TSSs in (A) mouse and (B) bovine. (C–H) Proportion of TE subfamilies containing specific transcription factor-binding motifs. The charts illustrate the distribution of (C) GATA3, (D) KLF5, and (E) OTX2 motifs within mouse TE sequences, and (F) DUXBL, (G) KLF5, and (H) OTX2 motifs within bovine TE sequences. Multiple matches of a single motif within a TE were included in the counts.

### Motif scanning on TEs shows motif-dependent enrichment with low cross-species conservation

Next, to examine whether the enriched transcription factor motifs were preferentially associated with specific TE subfamilies, we focused on GATA3, KLF5, and OTX2 in mouse and DUXBL, KLF5, and OTX2 in bovine; motif scanning was performed on those motifs across all TE sequences. As a result, in both mouse and bovine, almost all of these motifs were enriched in ERV superfamilies, including the ERVK family. The GATA3 binding motif was especially enriched in MT2_mm elements and in the RLTR family of ERVK elements (Fig. 5C). In both KLF5 and OTX2, the motifs were suggested to be located within the ORR1A family in mouse (Fig. 5D, 5E).

Interestingly, the top-ranked subfamilies for KLF5 motifs were LTR portions, whereas the motifs of OTX2 tended to be included in internal regions of LTR retrotransposons (designated “-int”) (Fig. 5D, E). In bovine, the DUXBL motif was found across both LINE and LTR retrotransposons, and its frequency of occurrence was lower than that of the other motifs, at less than 0.3 (Fig. 5F). The KLF5 and OTX2 motifs were both enriched mainly in the ERVK and ERV1 families in bovine. However, there was no tendency for these motifs to be located in specific TE portions, as observed in the ORR1A family in mouse, and the types of TEs enriched also differed from those in mouse (Fig. 5G, H).

## Discussion

Although chimeric transcripts generated by TE-derived promoters represent an important source of transcriptome diversity, their characterization using short-read sequencing has been hampered by the repetitive nature and abundance of TEs in the genome. Here, we constructed comprehensive catalogs of TE-driven chimeric transcripts expressed in mammalian embryos at the ZGA stage by applying nanopore long-read RNA sequencing, thereby suggesting the advantage of long-read sequencing for the analysis of chimeric transcripts.

In mouse, we identified 11,996 chimeric transcript variants from 2,695 host genes, exceeding the numbers reported in previous short-read-based studies[12], [13]. This result suggests that long-read sequencing is more advantageous for detecting chimeric transcripts. In addition, more than one-third of the genes identified in previous short-read-based studies overlapped with those identified in our study, indicating reasonable reproducibility despite differences in analytical methods.

We predicted ORFs for the detected TE-driven transcripts and analyzed their protein-coding potential. This analysis revealed an increased proportion of lncRNAs among TE-driven transcripts compared with the overall proportion among all detected transcripts. This observation is consistent with previous findings in mouse showing that lncRNAs contain a higher proportion of TE-derived sequences than protein-coding genes [35]. Notably, comparison of the ORFs from TE-driven transcripts classified as “Protein-coding” with their corresponding reference ORFs revealed a consistent pattern in both mouse and bovine: a substantial fraction of these transcripts exhibited truncated N termini while retaining their C-terminal regions. Although N termini can be more variable than C termini, because we focused on TE-driven genes, these findings suggest that the molecular mechanisms underlying the transcription and translation initiation of TE-driven transcripts may be conserved across mammalian species. Transcript variants harboring internal TE-derived exons have been shown to encode shorter proteins than their canonical counterparts in human cancer cells [36]. However, because the positions of TE-derived sequences differ between the transcripts identified in our study and those reported by Arribas et al., further investigation is required to elucidate the nature of the translational alterations in chimeric transcripts expressed in mammalian embryos. In cases where chimeric transcripts conserve the canonical ORF structure (with or without N-terminal truncation), TEs likely function as alternative promoters. The major ZGA stage in mouse embryos is preceded by a globally relaxed epigenetic landscape established during the minor ZGA stage, when complex transcriptional regulation is limited [37]. Under such conditions, TEs dispersed across the genome may serve as readily accessible regulatory elements [37], [38].

In addition, motif analysis of sequences upstream of TE-TSSs showed little conservation of enriched transcription factor motifs between species, except for KLF5 and OTX2. Moreover, the TEs harboring these motifs also differed: the KLF5 and OTX2 motifs were predominantly detected within mouse-specific LTR retrotransposon sequences in mouse but were distributed across more diverse TE families in bovine. These findings suggest that mouse-specific LTR retrotransposons may play a central role in transcriptional regulation during mouse embryogenesis, because they contain binding motifs for transcription factors active in mammalian embryos. In contrast, in bovine, the contribution of TEs to transcriptional regulation during embryogenesis appears to be lower than in mouse, with KLF5 and OTX2 binding motifs broadly distributed across diverse TSSs rather than associated with specific TEs. Furthermore, in mouse, KLF5 motifs were enriched in the LTR of ORR1A1, whereas OTX2 motifs were enriched within internal ORR1A1 sequences (ORR1A1-int), suggesting that distinct transcription factors may bind to different regions within the same TE family. Several transcription factors corresponding to the motifs identified in this analysis have previously been reported to be expressed after the major ZGA stage [30], [39], [40], [41]. Therefore, these transcription factors may contribute to transcriptional regulation at later developmental stages, rather than directly during ZGA.

Analysis of TE orders functioning as promoters revealed that LTR retrotransposons constituted a major fraction of TE-derived promoters in both species. However, at the subfamily level, mouse-specific TE subfamilies were enriched in mouse, whereas evolutionarily older subfamilies conserved across eutherian mammals were prominently represented in bovine. This finding suggests that TE-mediated transcriptional regulation is not strongly conserved across species but instead has evolved in a largely species-specific manner, with the dependence of transcriptional regulation on TEs may differ substantially between species. In mouse embryos, many studies have reported that mouse-specific LTR retrotransposons, such as MT2_Mm, are frequently utilized as promoters [42], [43]. In contrast, our study showed that evolutionarily older TEs predominate in bovine embryos at the ZGA stage. Indeed, the mouse-specific TE MT2B2 has been inserted upstream of *Cdk2ap1* and drives its transcription in mouse, whereas *CDK2AP1* is transcribed from the DNA transposon L2a/Charlie4 in many eutherian species [12]. A previous report suggested that bovine may have experienced fewer recent TE insertions during speciation compared with mouse [13] and, consequently, the turnover of TE-derived promoters in embryos may have been less extensive. Consistent with this, the number of TE-TSSs identified differed markedly between species, with approximately 8,000 in mouse and fewer than 3,000 in bovine. These findings suggest that transcriptional regulation in bovine embryos is less dependent on TEs than in mouse. Modzelewski et al. suggested that TEs contribute to species-specific gene expression and developmental processes, such as the rate of preimplantation development, thereby generating differences between species [12]. The differences observed here may likewise contribute to such developmental divergence.

## Conclusion

We performed long-read RNA sequencing of ZGA-stage embryos of mouse and bovine and comprehensively analyzed chimeric transcripts. We identified both the genes forming chimeric transcripts and the TE-derived promoters in greater numbers than in previous short-read sequencing studies, indicating an advantage of long-read sequencing for chimeric transcript analysis. However, while long-read sequencing enables the detection of full-length mRNAs, it is limited by lower accuracy and reduced quantitative reliability. Therefore, further validation will be required by integrating published datasets, including those for assessing histone modifications at the TSSs of identified chimeric transcripts, to add robustness to our datasets. The protein-coding potential and ORF characteristics of TE-driven transcripts were somewhat conserved across species, which suggests that the molecular mechanisms underlying transcription and translation from TEs share common features. In contrast, multiple analyses of TE promoters revealed marked differences between mouse and bovine, highlighting the species-specific nature of chimeric transcripts. These differences may reflect the promoter activity of TEs inserted during species divergence and could contribute to interspecies variation in developmental mechanisms. This study has limitations because only ZGA-stage embryos were analyzed. To achieve a comprehensive understanding of the contribution of chimeric transcripts during early embryogenesis, analyses across other developmental stages will be necessary.

## List of abbreviations

GO: Gene Ontology
HTF: Human tubal fluid
IVF: In vitro fertilization
KSOM: Potassium simplex optimized medium
LINE: Long interspersed nuclear elements
lncRNA: Long non-coding RNA
LTR: Long terminal repeat
ORF: Open reading frame
SINEs: Short interspersed nuclear elements
TE: Transposable element
TSS: Transcription start site
TE-TSS: TSSs that overlap TEs
ZGA: Zygotic genome activation

## Declarations

### Ethics approval and consent to participate

This study was conducted in accordance with the Regulation on Animal Experimentation at Kyoto University and approved by the Animal Experimentation Committee of Graduate School of Agriculture, Kyoto University (Approval Number: R5–10, R5–17, R6–10, and R6–17).

### Consent for publication

Not applicable.

### Availability of data and materials

The data of this study are available from the corresponding author upon request.

### Author’s contributions

S.H and S.K conceived and designed the study. S.K. collected the samples. S.K performed the morphological identification of the species. S.K performed experiments, including RNA extraction and library preparation and sequence. S.K performed the bioinformatic analyses and prepared all Figures with advice from K.K. S.K. wrote the main manuscript text. S.H supervised the project and critically revised the manuscript. All authors read and approved of the final manuscript.

---

**Supplementary Fig. 1.**
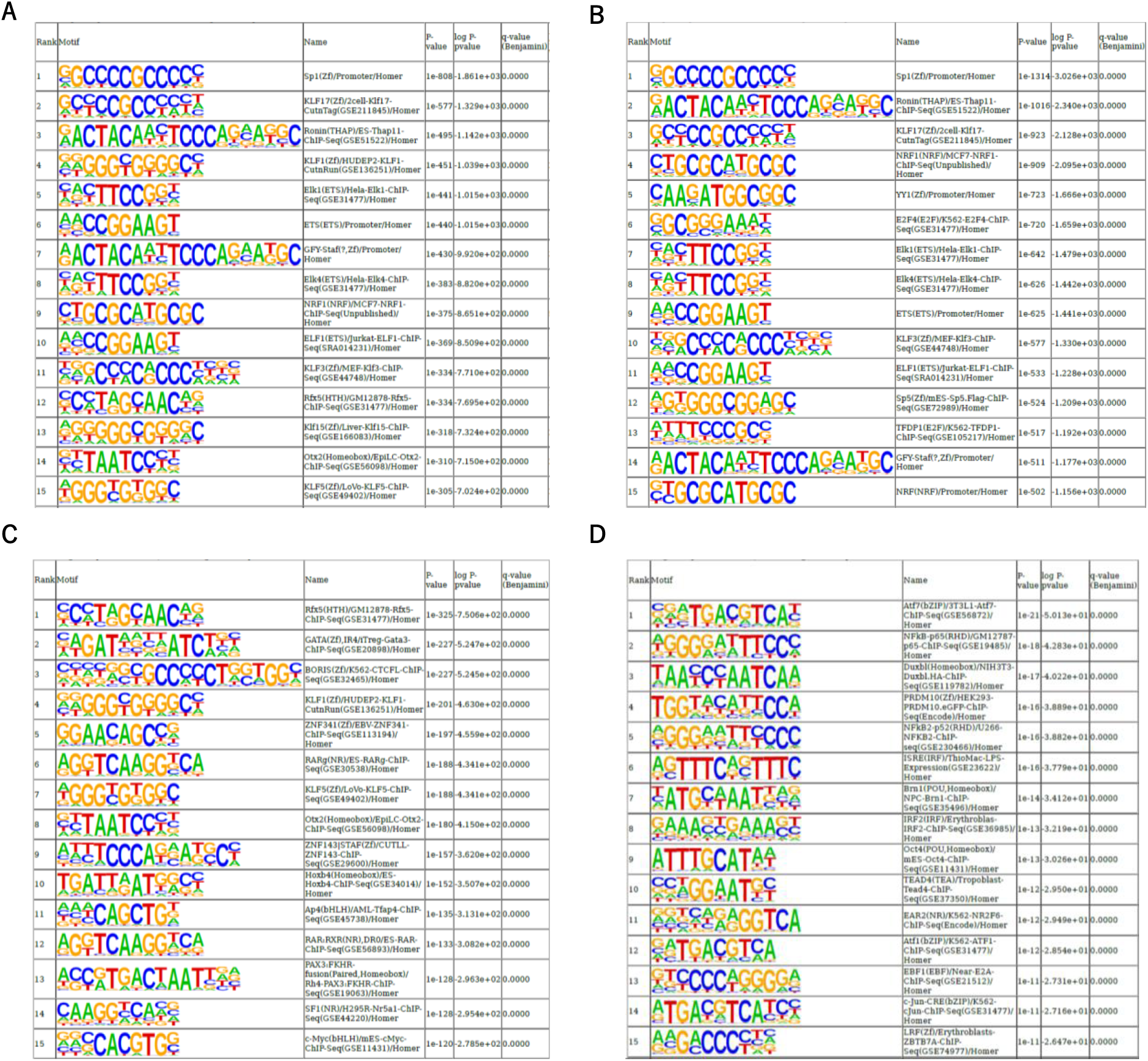
Motif analysis in the regions upstream of all TSSs and TE-TSSs. (A–B) The 15 most significantly enriched known motifs identified in regions within 1000 bp upstream of all identified TSSs in (A) mouse and (B) bovine. (C–D) The 15 most significantly enriched known motifs identified in regions within 1000 bp upstream of TE-TSSs in (A) mouse and (B) bovine, determined relative to a background set of all identified TSS sequences in each species.

**Table S1.**

| Sample | Primary Alignment (%) | Secondary Alignment | Supplementary alignment | Unmapped reads | Total Reads |
| --- | --- | --- | --- | --- | --- |
| Long_read (mouse) | 1562691 (100) | 0 | 0 | 0 | 1562691 |
| Long-read (bovine) | 2122809 (100) | 0 | 0 | 0 | 2122809 |

**Table S2.**

| Description | GeneRatio | BgRatio | pvalue | p.adjust | qvalue |
| --- | --- | --- | --- | --- | --- |
| DNA repair | 27/760 | 281/16137 | 0.000362 | 0.381842 | 0.368406 |
| meiotic cell cycle | 13/760 | 101/16137 | 0.000898 | 0.381842 | 0.368406 |
| hippo signaling | 6/760 | 27/16137 | 0.001353 | 0.381842 | 0.368406 |
| DNA metabolic process | 34/760 | 418/16137 | 0.001405 | 0.381842 | 0.368406 |
| glycerolipid metabolic process | 21/760 | 217/16137 | 0.001433 | 0.381842 | 0.368406 |
| meiotic cell cycle process | 12/760 | 94/16137 | 0.001502 | 0.381842 | 0.368406 |
| DNA damage response | 30/760 | 356/16137 | 0.001527 | 0.381842 | 0.368406 |
| nucleic acid catabolic process | 14/760 | 126/16137 | 0.002439 | 0.434297 | 0.419015 |
| muscle cell development | 7/760 | 41/16137 | 0.00277 | 0.434297 | 0.419015 |
| striated muscle tissue development | 6/760 | 31/16137 | 0.002868 | 0.434297 | 0.419015 |

B
| Description | GeneRatio | BgRatio | pvalue | p.adjust | qvalue |
| --- | --- | --- | --- | --- | --- |
| protein serine/threonine kinase activity | 31/731 | 321/16210 | 5.66E-05 | 0.033338 | 0.03253 |
| sodium ion transmembrane transporter activity | 13/731 | 117/16210 | 0.002362 | 0.420734 | 0.410546 |
| SMAD binding | 5/731 | 22/16210 | 0.002553 | 0.420734 | 0.410546 |
| gamma-tubulin binding | 4/731 | 14/16210 | 0.002857 | 0.420734 | 0.410546 |
| catalytic activity, acting on a nucleic acid | 33/731 | 467/16210 | 0.007336 | 0.625276 | 0.610134 |
| I-SMAD binding | 3/731 | 10/16210 | 0.008638 | 0.625276 | 0.610134 |
| catalytic activity, acting on DNA | 16/731 | 183/16210 | 0.008742 | 0.625276 | 0.610134 |
| molecular sensor activity | 4/731 | 19/16210 | 0.009248 | 0.625276 | 0.610134 |
| phosphatidylinositol kinase activity | 4/731 | 20/16210 | 0.011155 | 0.625276 | 0.610134 |
| endodeoxyribonuclease activity, producing 5'-phosphomonoesters | 3/731 | 11/16210 | 0.011484 | 0.625276 | 0.610134 |

C
| Description | GeneRatio | BgRatio | pvalue | p.adjust | qvalue |
| --- | --- | --- | --- | --- | --- |
| DNA recombination | 9/113 | 343/28832 | 8.67E-06 | 0.018536 | 0.015626 |
| protein autophosphorylation | 6/113 | 200/28832 | 0.000141 | 0.075828 | 0.063922 |
| positive regulation of dendritic spine development | 4/113 | 68/28832 | 0.00015 | 0.075828 | 0.063922 |
| dendritic spine development | 5/113 | 139/28832 | 0.000224 | 0.075828 | 0.063922 |
| regulation of mRNA metabolic process | 7/113 | 321/28832 | 0.00028 | 0.075828 | 0.063922 |
| execution phase of apoptosis | 4/113 | 80/28832 | 0.000281 | 0.075828 | 0.063922 |
| prostaglandin biosynthetic process | 3/113 | 33/28832 | 0.000294 | 0.075828 | 0.063922 |
| prostanoid biosynthetic process | 3/113 | 33/28832 | 0.000294 | 0.075828 | 0.063922 |
| chromosome segregation | 8/113 | 434/28832 | 0.000319 | 0.075828 | 0.063922 |
| regulation of dendritic spine development | 4/113 | 88/28832 | 0.000405 | 0.08659 | 0.072995 |

D
| Description | GeneRatio | BgRatio | pvalue | p.adjust | qvalue |
| --- | --- | --- | --- | --- | --- |
| histone H2AX kinase activity | 11/113 | 359/28366 | 2.11E-07 | 1.11E-05 | 8.96E-06 |
| protein serine kinase activity | 11/113 | 362/28366 | 2.29E-07 | 1.11E-05 | 8.96E-06 |
| histone H3 kinase activity | 11/113 | 362/28366 | 2.29E-07 | 1.11E-05 | 8.96E-06 |
| histone kinase activity | 11/113 | 366/28366 | 2.55E-07 | 1.11E-05 | 8.96E-06 |
| 3-phosphoinositide-dependent protein kinase activity | 9/113 | 250/28366 | 7.48E-07 | 1.11E-05 | 8.96E-06 |
| DNA-dependent protein kinase activity | 9/113 | 250/28366 | 7.48E-07 | 1.11E-05 | 8.96E-06 |
| eukaryotic translation initiation factor 2alpha kinase activity | 9/113 | 250/28366 | 7.48E-07 | 1.11E-05 | 8.96E-06 |
| ribosomal protein S6 kinase activity | 9/113 | 250/28366 | 7.48E-07 | 1.11E-05 | 8.96E-06 |
| histone H3S10 kinase activity | 9/113 | 250/28366 | 7.48E-07 | 1.11E-05 | 8.96E-06 |
| histone H2AXS139 kinase activity | 9/113 | 250/28366 | 7.48E-07 | 1.11E-05 | 8.96E-06 |

